# Effects of Post-Myocardial Infarction Heart Failure on the Bone Vascular Niche

**DOI:** 10.1101/2020.05.29.123711

**Authors:** Jedrzej Hoffmann, Guillermo Luxán, Wesley Tyler Abplanalp, Simone-Franziska Glaser, Tina Rasper, Ariane Fischer, Marion Muhly-Reinholz, David John, Birgit Assmus, Andreas Michael Zeiher, Stefanie Dimmeler

## Abstract

Bone vasculature provides protection and signals necessary to control stem cell quiescence and renewal^1^. Specifically, type H capillaries, which highly express Endomucin, constitute the endothelial niche supporting a microenvironment of osteoprogenitors and long-term hematopoietic stem cells^2–4^. The age-dependent decline in type H endothelial cells was shown to be associated with bone dysregulation and accumulation of hematopoietic stem cells, which display cell-intrinsic alterations and reduced functionality^3^. The regulation of bone vasculature by chronic diseases, such as heart failure is unknown. Here, we describe the effects of myocardial infarction and post-infarction heart failure on the vascular bone cell composition. We demonstrate an age-independent loss of type H bone endothelium in heart failure after myocardial infarction in both mice and in humans. Using single-cell RNA sequencing, we delineate the transcriptional heterogeneity of human bone marrow endothelium showing increased expression of inflammatory genes, including *IL1B* and *MYC*, in ischemic heart failure. Inhibition of NLRP3-dependent IL-1β production partially prevents the post-myocardial infarction loss of type H vasculature in mice. These results provide a rationale for using anti-inflammatory therapies to prevent or reverse the deterioration of vascular bone function in ischemic heart disease.

## Main

Recent characterization of endothelial cells (EC) in the murine bone led to the identification of at least two functional vessel subsets, based on their differential high (H) and low (L) expression of Endomucin (EMCN) and CD31. Type H capillaries are arteriole-associated, columnar vessels in the metaphysis and endosteum regions. Type L vessels are sinusoid-associated vessels, which predominate the whole medullary region^1^. Age-dependent decline of type H endothelium is associated with bone dysregulation and accumulation of long-term haematopoietic HSCs (LT-HSC)^3^. Although the functional interaction between heart and bone is emerging as a trigger of post-infarction inflammation and progression of cardiovascular disease^5^, the impact of myocardial infarction (MI) and resulting heart failure on the vascular niche in the bone remains unknown. Therefore, we induced MI in mice (12 week-old) by left anterior descending (LAD) coronary artery ligation^6^ (Supplementary Fig. 1) and assessed the kinetics of bone EC subsets, haematopoietic progenitor, and stem cells during the development of post-MI heart failure. MI induced a time-dependent steady reduction of type H EC frequencies as assessed by flow cytometric analysis. Type H cells were significantly decreased at day 7 after infarction and profoundly diminished by day 28 as compared to control mice (Fig. 1a, b). This observation was confirmed by histological analysis of mouse femurs (Fig. 1c). Coinciding with the decrease of type H endothelium, LT-HSCs significantly increased in mice during development of post-infarction heart failure (Fig. 1b and Supplementary Fig. 2a). The reduction in type H vessels was not age dependent, as histological assessment confirmed that type H vessel length was significantly reduced 28 days after infarction when compared to age matched controls (Supplementary Fig. 2b). Detailed analysis of CD41 immunostainings of the femur marrow, which is a hallmark of impaired vascular niche function, revealed a significant increase in CD41 positive cells, suggesting a myeloid bias in the mice that underwent LAD ligation^7^ (Fig. 1d).

**Figure 1.**
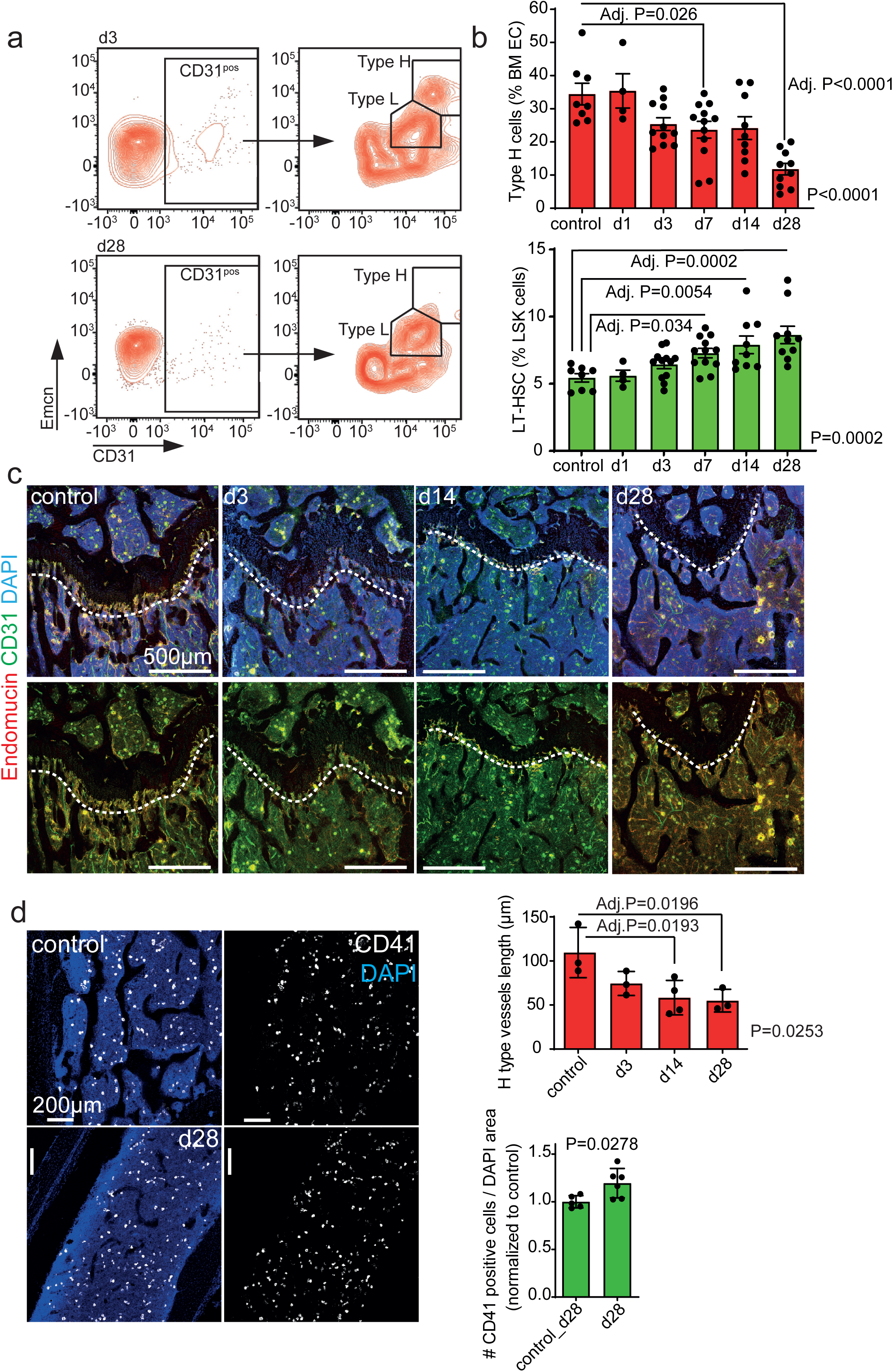
Type H endothelium is reduced in the femur upon myocardial infarction (MI). (**a, b**) Flow cytometry analysis of femur bone marrow. Surgery was performed on 12-week-old animals. (**a**) Gating strategy for endothelial cells. Representative samples at day (d) 3 and d28 are shown. (**b**) Quantification. Upper panel, type H endothelial cells are reduced after MI. Lower panel, LT-HSC cells are increased after MI. N=8 for control, N=4 for d1, N=12 for d3, N=12 for d7, N=9 for d14 and N=10 for d28. Data are shown as mean ± SEM. P-value was calculated with ANOVA with Dunnet’s multiple comparison test. (**c, d**) Immunostaining of longitudinal sections through femur. (**c**) The length of type H vessels (indicated by the dashed line) is reduced after MI. N=3 for control, d3 and d28 and N=4 for d14. Data are shown as mean ± SEM. P-value was calculated with ANOVA. Comparisons between all groups was calculated by Dunnet’s multiple comparison test. (**d**) Myeloid progenitor cell number is increased in the bone marrow 28 days after MI. N=5 for each condition. Data are shown as mean ± SEM. P-value was calculated by unpaired two-tailed Student’s *t*-test.

To confirm the relevance of these findings in humans, we determined the number and expression pattern of endothelial cells in bone marrow aspirates of healthy volunteers and patients with post-MI heart failure (for baseline characteristics of patients see Supplementary Table 1). Flow cytometry analysis showed that elderly heart failure patients’ display significantly reduced numbers of CD31^hi^EMCN^hi^ representing type H endothelial cells in the bone marrow aspirates, whereas total bone marrow endothelial cells were not significantly altered (Figure 2a, b). To gain deeper insights into the age-independent regulation of the vascular niche by heart failure, we performed single-cell RNA sequencing of lineage-depleted CD31 enriched cells obtained from the bone marrow aspirates of an age-matched healthy volunteer and a patient with post-MI heart failure (Supplementary Table 1). t-Stochastic-Neighbour-Embedding (t-SNE) analysis revealed 13 clusters with equal distribution of cells from the healthy donor and the heart failure patient among the clusters (Fig. 2c). Annotating of the clusters (Fig. 2c middle panel) showed that cells expressing *EMCN* transcript were significantly enriched in cluster 0, representing the type H endothelial cell population (Fig. 2c, d and Supplementary Fig. 3). Re-clustering of these cells revealed a striking divergence in the populations of cells derived from the heart failure patient and the age matched healthy control (Fig 2e, left). Analysis of differentially expressed genes between these two new populations did not reveal differences in the expression of endothelial genes like *PECAM1 (CD31)* (Fig. 2e- right, f). However, inflammatory gene transcripts, specifically *IL1B*^8,9^ and *MYC*^10^ transcripts, were significantly increased in the heart-failure bone marrow sample (Fig 2e-right, f). The increase in *IL1B* and *MYC*, was confirmed at protein level by representative immunocytochemistry analysis of bone marrow endothelial cells (Supplementary Fig. 4). Pseudotime trajectory analysis of re-clustered *EMCN, PECAM1* enriched cells revealed 6 branch points and 13 different states (Fig. 2g), four of which (states 10 to 13) were predominately populated by the cells derived from the heart failure patient (Fig. 2g). These states expressed higher levels of *IL1B* (Fig. 2h). Upregulated genes from state 13 (heart failure enriched) revealed activation of Wnt and NF-κB signalling pathways (Fig. 2i).

**Figure 2.**
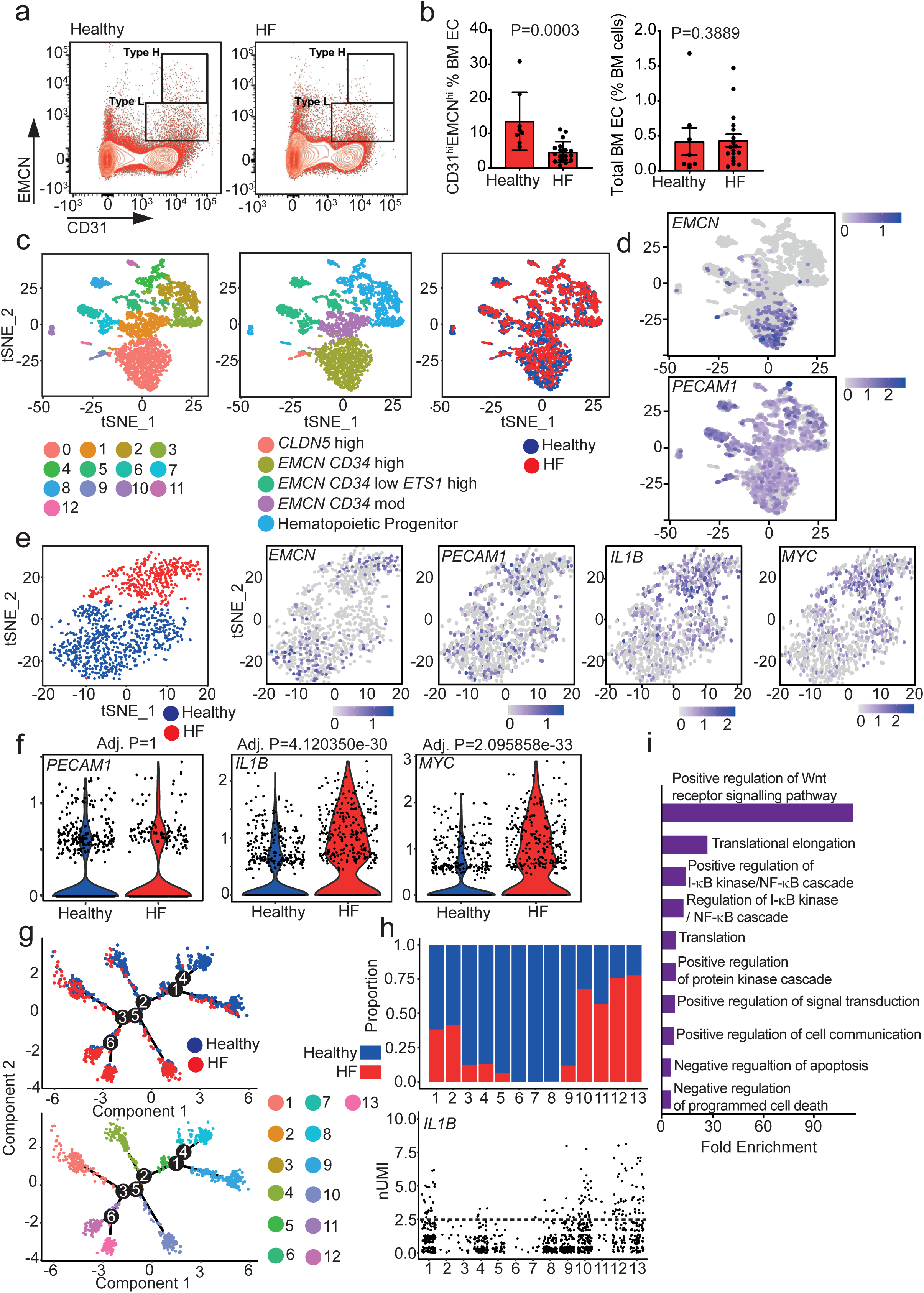
*IL1B* is upregulated in *EMCN* rich endothelial cells in post-myocardial infarction (MI) heart failure (HF). (**a, b**) Flow cytometry analysis of healthy and HF patient bone marrow. (**a**) Gating strategy for endothelial cells. Representative flow cytometry dot plots showing EC subsets with distinct expression of CD31 and Endomucin (EMCN) in healthy and HF bone marrow aspirates (gated on CD45^**neg**^Lin^**neg**^ viable single cells). (**b**) Type H endothelial cells are reduced in the HF patients compared to healthy controls (left panel), while the total number of endothelial cells remains unchanged (right panel) (N=8 for healthy, N=18 for HF patients). Data are shown as mean ± SEM. P-value was calculated by unpaired, Mann Whitney test. (**c**-**i**) scRNA-seq of a post-MI heart failure and an aged-matched healthy control. (**c**,**d**) Clustered cells from both subjects are displayed in t-SNE plots, colored by cluster (left), cell annotation (middle) and health status (right). (**d**) Expression of *EMCN* and *PECAM1. EMCN* is enriched in the cells corresponding to cluster 0. (**e**-**f**) Analysis of *EMCN* enriched cell cluster 0 population. (**e**) Dichotomization in *EMCN* enriched population shown in t-SNE plot (Left). Relative expression of key genes in the *EMCN* enriched population represented by features plots as indicated. (**f**) Violin plots showing the relative expression of key genes in the *EMCN* enriched population, confirming the significantly increased expression of *IL1B* and *MYC* in the HF patient. (**g**) Distribution of cells along pseudotime trajectory branchpoints. Pseudotime analysis revealed 13 states. (**h**) Distribution of cells among pseudotime states and relative *IL1B* expression. Distribution analysis revealed that states 10, 11, 12, and 13 are mainly populated by HF patient cells. *IL1B* expression is higher in states 12 and 13. Dashed line indicates normalized Unique Molecular Identifier (nUMI) counts of 2.5. (**i**). Gene Ontology term ranking of upregulated genes in pseudotime state 13.

Interleukin-1β is a prototypic proinflammatory cytokine, processed to its active form in different cell types mostly by activated inflammasome complexes, such as NLRP3^11^. Therefore, we further characterised the kinetics of IL-1β in bone endothelial cells and the impact of its expression on the vascular niche phenotype in post-infarction heart failure. First, we investigated the global *Il1b* expression in the bones from mice after infarction. RT-qPCR from total bone marrow of mice after LAD ligation confirmed a steady increase in expression of *Il1b* during the development of heart failure, with highest expression at day 28 post-infarction (Fig. 3a). To gain further insights into the specific expression of IL-1β in the bone marrow vasculature, we performed immunostainings of bone sections after MI. Strikingly, there was a strong induction of IL-1β, specifically in type H ECs, as early as 1 day after induction of MI (Fig. 3b). This rapid induction of IL-1β in type H endothelium preceded the subsequent accelerated loss of type H cells in the course of MI.

**Figure 3.**
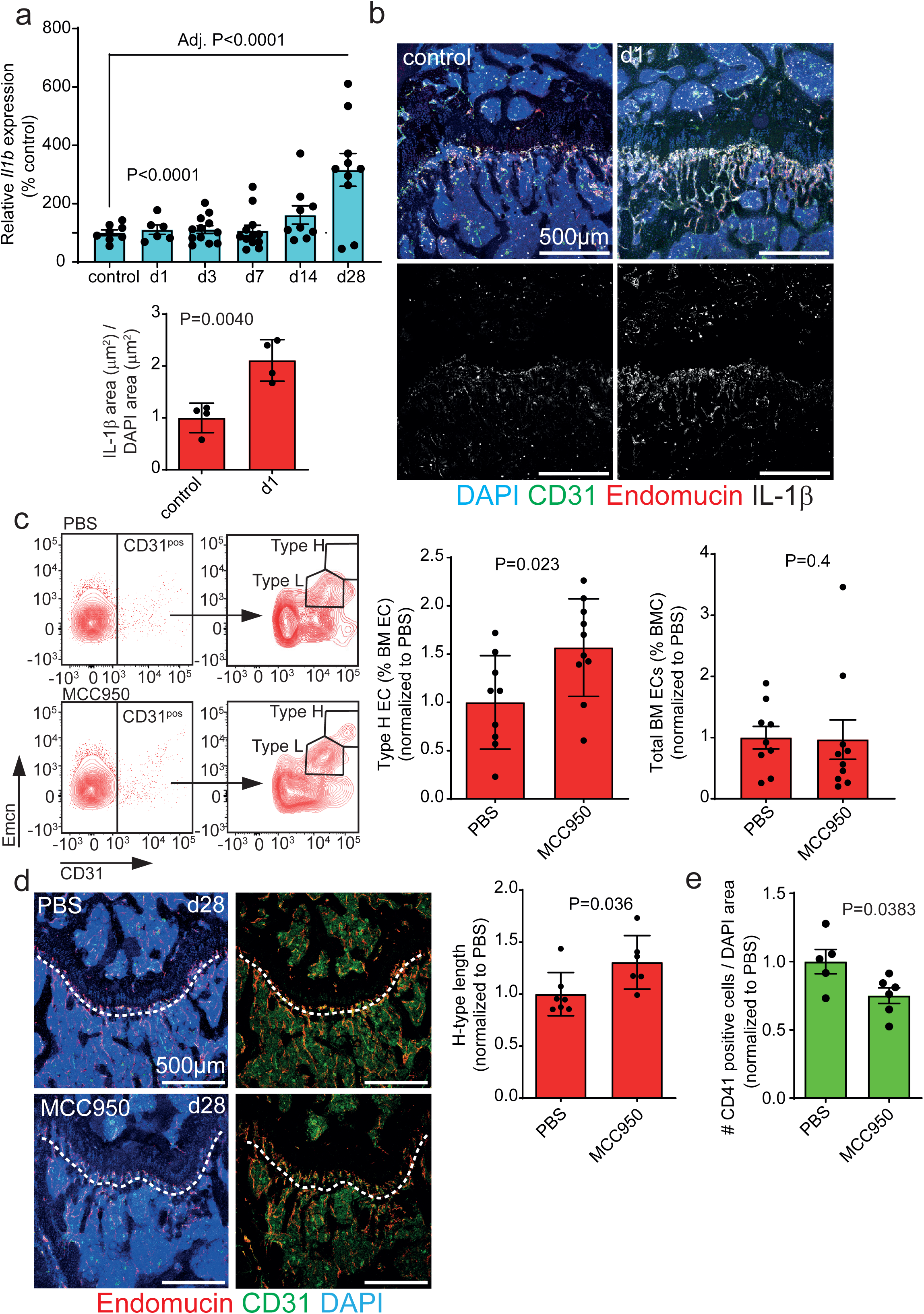
Anti-IL-1β treatment ameliorates the loss of type H vasculature in the bone after MI. (**a**) RT-qPCR from total femur bone marrow. MI induces *Il1b* expression in the bone marrow. N=8 for Control, N=6 for d1, N=12 for d3, N=12 for d7, N=9 for d14 and N=10 for d28. Data are shown as mean ± SEM. P-value was calculated with ANOVA. Comparisons between all groups was calculated by Dunnet’s multiple comparison test. (**b**) Immunostaining of longitudinal sections through femur. IL-1β immuno-signal is concentrated on type H vessels one day after infarct. N=4 for each condition. Data are shown as mean ± SEM P-value was calculated by unpaired, two-tailed Student’s t test. (**c**) Flow cytometry analysis of femur bone marrow. Left, gating strategy; right, quantification. Type H endothelial cell number is increased upon anti-IL-1β treatment, while total number of endothelial cells is not affected. N=9 for PBS and N=10 for MCC950. Data are shown as mean ± SEM. P-value was calculated by unpaired, two-tailed Student’s *t*-test (type H endothelial cell number) and Mann-Whitney test (total number of endothelial cells). (**d**) Immunostaining of longitudinal sections through femur. The length of type H vessels (dashed line) is longer upon MCC950 treatment 28 days after MI. N=7 for PBS and N=6 for MCC950. Data are shown as mean ± SEM. P-value was calculated by unpaired, two-tailed Student’s *t*-test. (**e**) Quantification of immunostaining in Supplementary Figure 5c. 3. Myeloid progenitor cell number is decreased in the bone marrow of MCC950 treated animals after infarct. N=5 PBS and N=6 MCC950. Data are shown as mean ± SEM. P-value was calculated by unpaired, two-tailed Student’s *t*-test.

Since our data suggest that MI alters the inflammatory environment of the vascular niche and particularly augments IL-1β expression, we determined whether IL-1β might play a causal role in mediating the deterioration of the vascular niche in post-MI heart failure. Therefore, we blocked IL-1β production with the selective NLRP3 inflammasome inhibitor MCC950^12^. MCC950 treatment partially prevented the MI- induced loss of type H endothelial cells in the bone compared to PBS treated controls as shown by flow cytometry (Fig. 3c) and bone section immunostainings (Figure 3d). MCC950 treatment did not have significant effects on the number of LT-HSC (Supplementary Fig. 5a, b), but reduced the number of CD41 positive cells (Fig. 3e and Supplementary Fig. 5c).

Here, we show for the first time an impact of post-MI heart failure on the bone marrow vasculature in both, mice and humans. The reduction of type H bone endothelial cells seems to be strongly associated with inflammatory responses, as evidenced by the induction of IL-1β in type H vessels, preceding their loss after MI. The question what directly triggers the observed strong IL-1β induction in type H cells after MI remains open. Regarding the observed NLRP3-inflammasome-dependent partial rescue of the vascular phenotype, one may consider a potential role of circulating damage-associated molecular patterns (DAMPs) being released from acute and chronically injured cardiac tissue. DAMPs would stimulate NLRP3 activation in endothelial and immune cells, leading to primary and secondary (by e.g. systemic and paracrine cytokines) IL-1β induction, release and amplification of inflammatory circuits^11^. Another possible mechanism comprises acute β-adrenergic stimulation at the bone level by adrenergic nerves in the course of MI. Increased β2- adrenergic activity has been shown to substantially influence niche microenvironment by promoting IL-6-dependent CD41^+^ myeloid progenitor expansion^13^. Inflammatory activation of specifically CD31^hi^EMCN^hi^ endothelial cells would result in pyroptotic or apoptotic cell death^14^. This might trigger an age-independent disruption of endothelium instructive function in the bone, leading to dysregulation of homeostatic haematopoiesis and HSC activity, marked by expansion and myeloid lineage skewing of HSC^15,16^. Our findings demonstrating that inhibition of IL-1β partially reversed the reduction of type H cells in the vascular niche and inhibited CD41^+^ myeloid progenitor cell expansion after MI provides mechanistic support for the therapeutic benefits of the anti-IL-1β antibody canakinumab. This therapy was shown, in a large clinical trial not only to reduce cardiovascular events in patients with previous MI (CANTOS trial)^17^, but - more importantly – inhibited the progression of post-MI heart failure^18^. Therefore, inhibition of the effector cytokine IL-1β, e.g. using canakinumab, might be considered as a novel strategy to prevent or reverse the deterioration of the vascular bone function in ischemic heart disease.

Furthermore, as type H endothelial cells maintain perivascular osteoprogenitors and couple angiogenesis to osteogenesis, their loss would also affect the bone forming cells and in consequence bone formation and maintenance. This proposed mechanism supports our previous observations^19^ and could help to further disclose a pathophysiological pathway linking post-MI heart failure with bone remodelling associated with an increased osteoporotic fracture risk^20,21^.

## Acknowledgements

We thank Eva-Maria Rogg and Marga Müller-Ardogan for excellent technical support and patient care and Halvard Bönig for support in establishing and validation of human flow cytometry panels.

## Author contribution

J.H., Conceptualization, Data curation, Formal analysis, Investigation, Funding acquisition, Writing—original draft. G.L., Conceptualization, Data curation, Formal analysis, Investigation, Writing—original draft. W.T.A., Data curation, Formal analysis, Investigation. S.F.G. Investigation, Methodology. T.R., Investigation, Methodology. A.F., Investigation, Methodology. M.M.R., Investigation, Methodology. D.J., Investigation, Methodology. B.A., Investigation, Methodology, Patient recruitment, Funding acquisition. A.M.Z., Conceptualization, Supervision, Funding acquisition, Project administration. S.D., Conceptualization, Supervision, Funding acquisition, Writing—original draft, Project administration

None of the authors have competing interests to declare

---

**Supplementary Figure 1.**
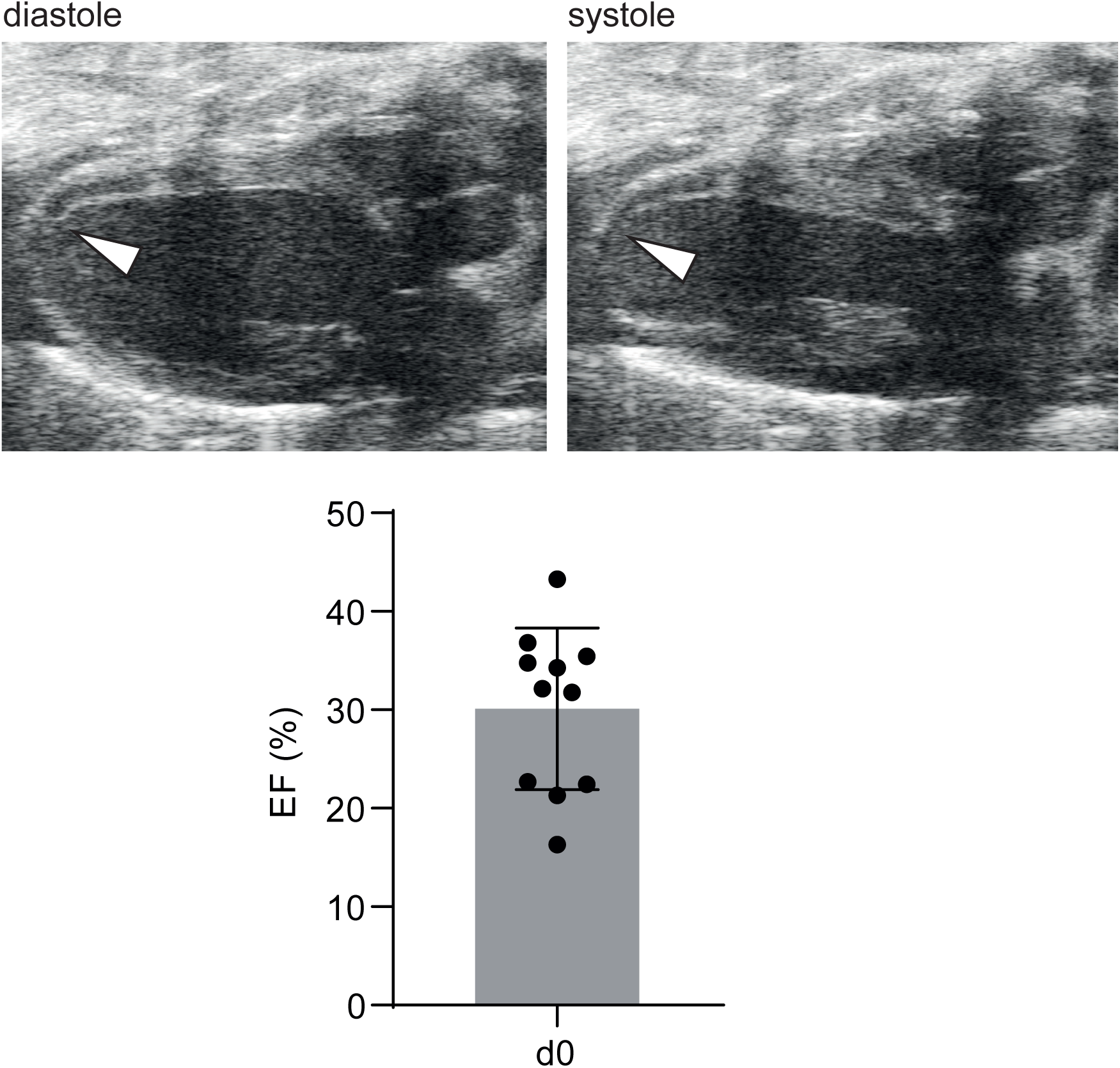
Representative echocardiographic analysis of animals after induced myocardial infarction. Arrowhead indicates the infarcted wall of the left ventricle. Lower panel, quantification of EF after MI.

**Supplementary Figure 2.**
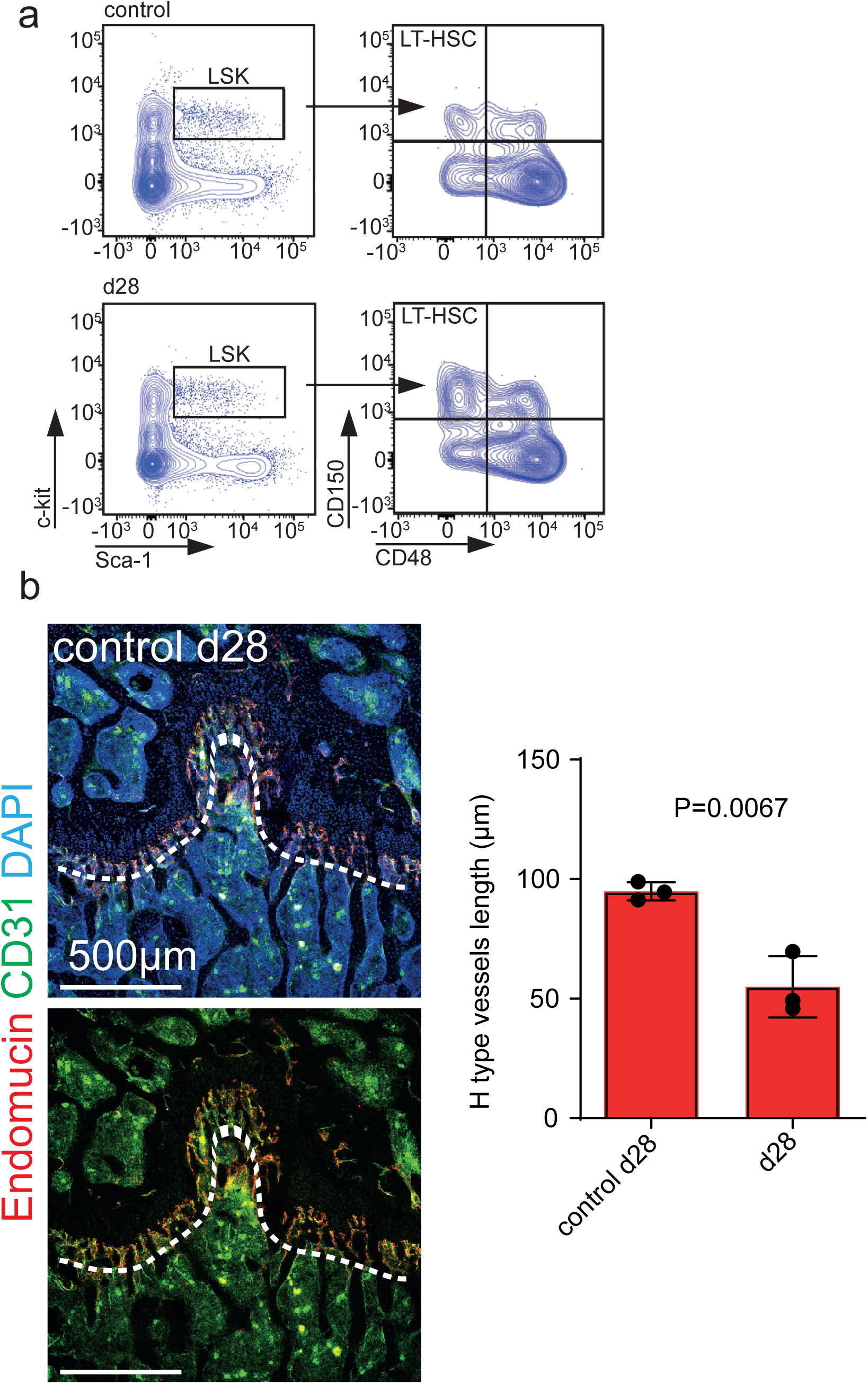
Type H reduction upon MI is independent of age. (**a**) Flow cytometry analysis of femur bone marrow. Surgery was performed on 12-week-old animals. Gating strategy for hematopoietic stem cells. (**b**) Immunostaining of longitudinal sections through age matched d28 control femur. Type H vessels are longer in the age matched d28 control than in mice 28 days after MI. N=3 for control d28. Data are shown as mean ± SEM. P-value was calculated by unpaired, two-tailed Student’s *t*-test.

**Supplementary Figure 3.**
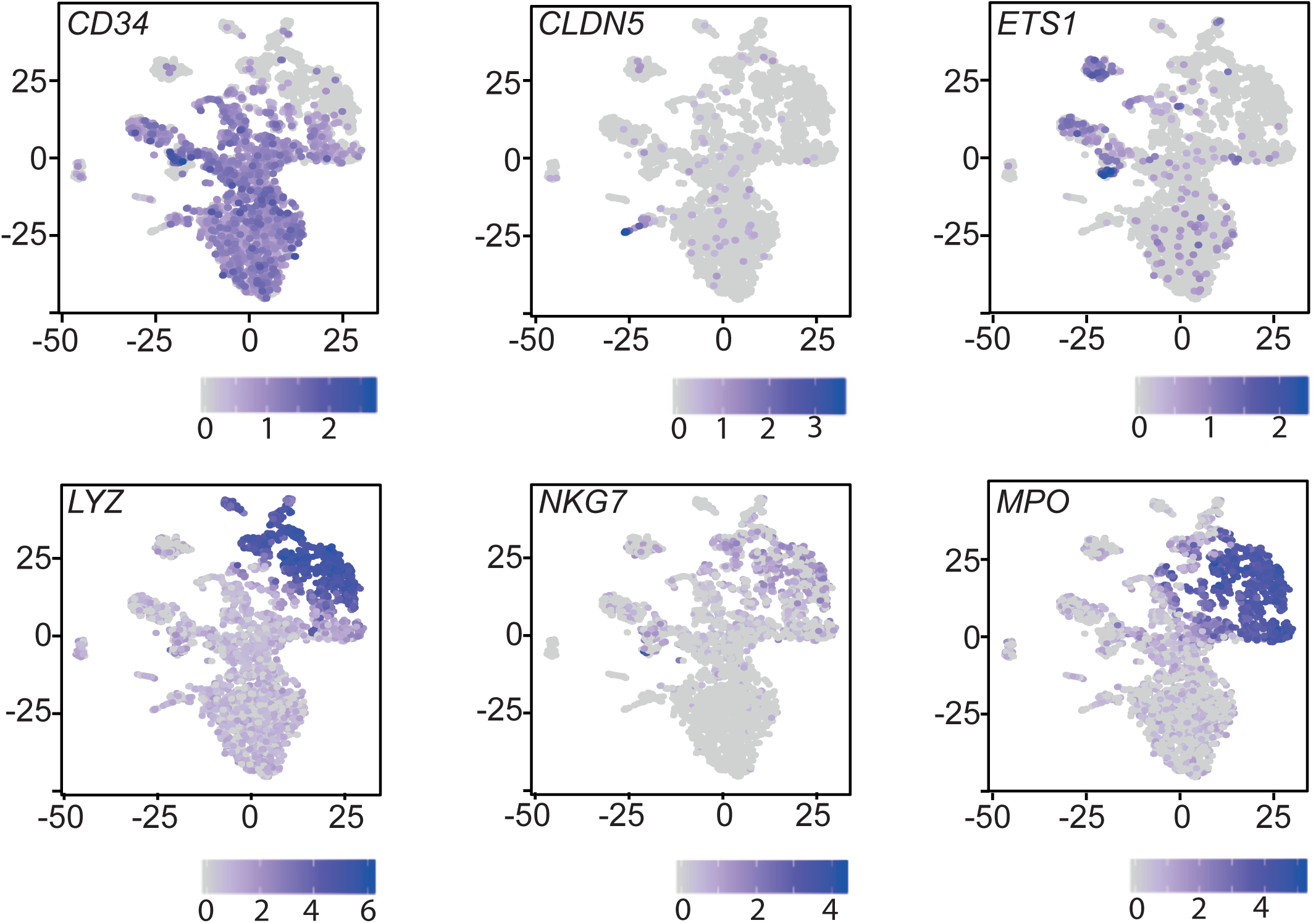
Expression of hallmark genes from scRNA-seq of bone marrow aspirates.

**Supplementary Figure 4.**
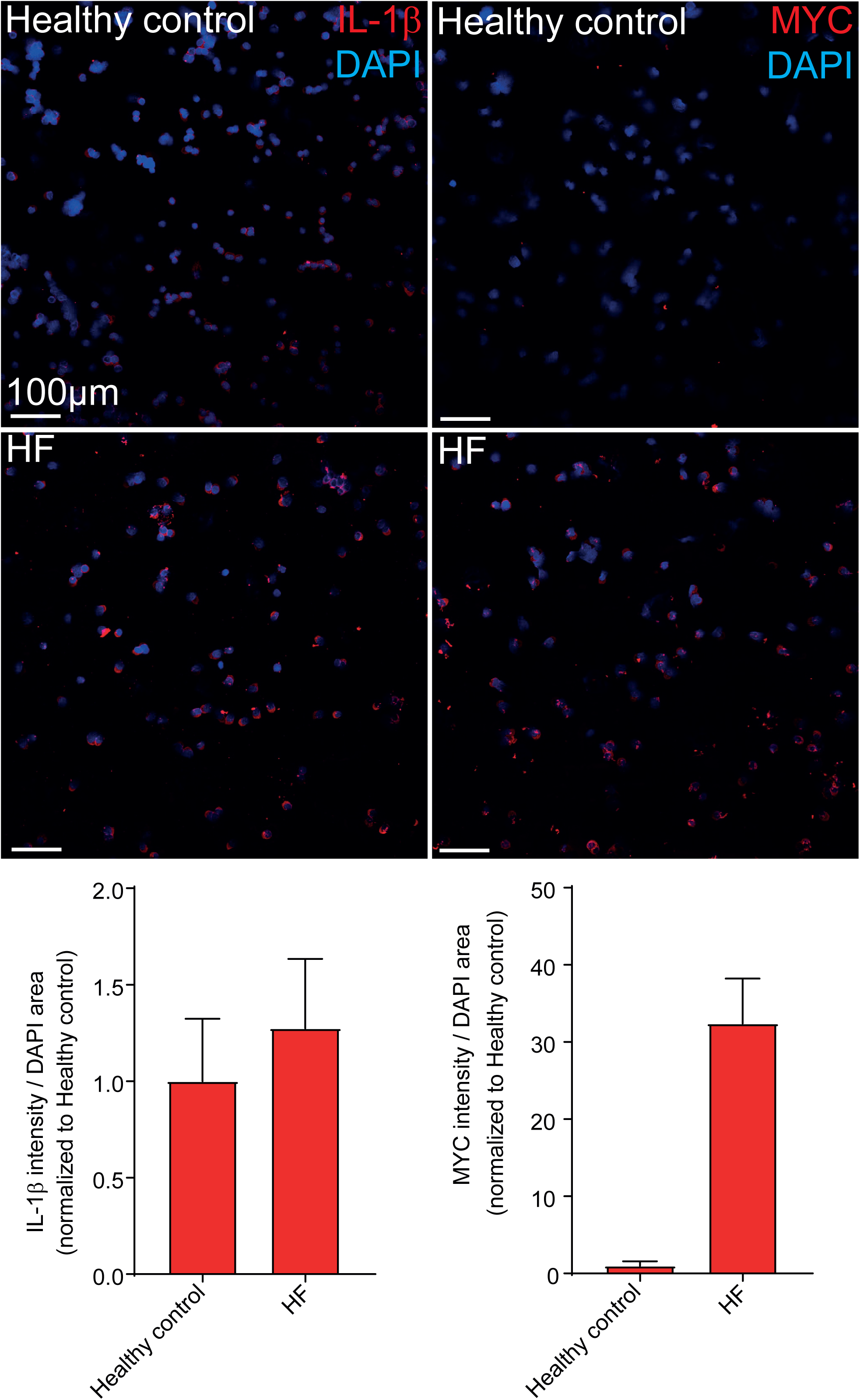
MYC and IL-1β are increased on patient bone marrow. Representative Immunostaining of MACS-enriched endothelial cells from freshly bone marrow aspirates. The expression of MYC and IL-1β is increased in the endothelial cells obtained from the post-ischemic heart failure patient bone marrow. Data are shown as mean ± SEM of the individual technical replicates per patient.

**Supplementary Figure 5.**
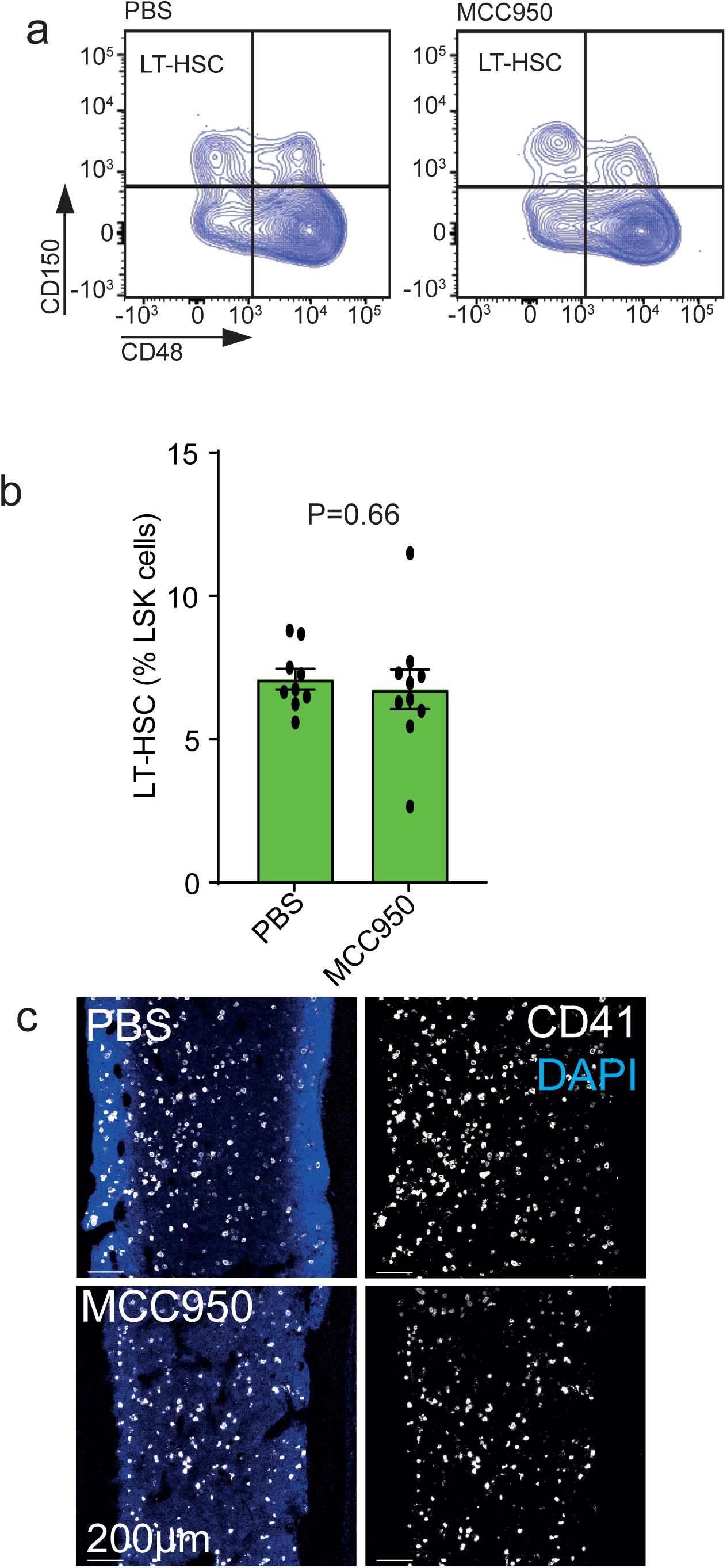
Anti-IL-1β treatment reduces myeloid progenitor number in the bone marrow. (**a, b**) Flow cytometry analysis of femur bone marrow. Surgery was performed on 12-week-old animals. (**a**) Gating strategy for hematopoietic stem cells. (**b**) Total number of LT-HSC remains unchanged after IL- 1β treatment. N=9 PBS and N=10 MCC950. Data are shown as mean ± SEM. P- value was calculated by unpaired, two-tailed Student’s *t*-test. (**c**) Immunostaining of longitudinal sections through femur. Myeloid progenitor cell number is decreased in the bone marrow of MCC950 treated animals after infarct.

**Supplementary Table 1.** Baseline characteristics of the post-MI heart failure patient study cohort

|  | Post-MI HF (FACS) | Post-MI HF (scRNA-Seq) |
| --- | --- | --- |
| n | 18 | 1 |
| Age (years) | 60 [IQR 54-71] | 43 |
| Sex (male/female) | 17/1 | 1/0 |
| NYHA class [I/II/III/IV (n)] | 0/9/9/0 | 0/1/0/0 |
| Hypertension [n (%)] | 13 (72) | 1 |
| Hypercholesterolemia [(%)] | 12 (67) | 1 |
| Diabetes [n (%)] | 4 (22) | 0 |
| Smoking [n (%)] | 14 (78) | 1 |
| Coronary artery disease<br>[1/2/3 vessel disease (n)] | 4/6/8 | 0/1/0 |
| Time from last MI (months) | 60 [IQR 15-145] | 7 |
| Left ventricular EF (%) | 28 [IQR 22-35] | 23 |
| ICD or CRT-D (%) | 15 (83) | 1 |
| ACEi or ATRB (%) | 17 (94) | 1 |
| Beta-blocker (%) | 16 (89) | 1 |
| Diuretic (%) | 16 (89) | 1 |
| Aldosterone antagonist | 15 (83) | 1 |
| Statin | 18 (100) | 1 |

HF- heart failure
MI – myocardial Infarction
EF – ejection fraction
ICD – implantable cardioverter defibrillator
ACEi – Angiotensin-converting enzyme inhibitor
ATRB - Angiotensin II receptor blocker

## Material and Methods

### Mouse strains

C57Bl/6J mice were used in this study. All animal procedures were performed according to relevant laws and institutional guidelines, were approved by local animal ethics committees and were conducted with permissions FU/1218 and FU/1222 granted by the Regierungspräsidium Darmstadt of Hessen.

### Myocardial Infarction

Myocardial infarction was performed in 12-weeks-old male C57Bl/6J mice. Under mechanical ventilation, the acute myocardial infarction (AMI) was induced by permanent ligation of the left anterior descending coronary artery.

### *In vivo* inhibition of the NLRP3 inflammasome

MCC950, a small molecule inhibitor of the NLRP3 inflammasome (S7809, Selleck Chemicals, Houston, USA) diluted in PBS and delivered *in vivo* at a dose of 5 mg/kg/day via subcutaneous mini-osmotic pumps (1004, ALZET), starting at the timepoint of AMI induction. Control mice were infused with PBS.

### Mouse flow cytometry

#### Isolation of mouse bone marrow

For flow cytometric analysis, tibiae and femurs were collected, cleaned thoroughly to remove the adherent muscles. Cleaned bones were then crushed in ice cold PBS with mortar and pestle. Whole bone marrow was digested with collagenase II (420U, Millipore) incubation at 37°C for 20 minutes. After digestion, cells were washed two times with PBS and filtered through a 100 µm cell strainer (EASYstrainer™ 100 µm, Greiner Bio-One). The cell concentration was adjusted to 10^6^/100µl in cell stain buffer (Biolegend).

#### HSC panel

Equal number of cells were then blocked with 2µl FC-Block (Mouse BD Fc Block™) per 100µl cell suspension for 10 minutes. After blocking cells were stained with FITC-conjugated Lineage cocktail (133301, Biolegend, clones 145- 2C11, RB6-8C5, M1/70, RA3-6B2, Ter-119), BV421- conjugated c-kit (105827, Biolegend, clone 2B8), PE-Cy7-conjugated Ly-6A (122513, Biolegend, clone E13- 161.7), APC-conjugated CD48 (103411, Biolegend, clone HM48-1) and PE-conjugated CD150 (115903, Biolegend, clone TC15-12F12.2) for 30 minutes on ice. After washing and adding 7AAD viability staining solution (420404, Biolegend), cells were acquired on BD FACS Canto II flow cytometer and analysed using FlowJo (Version 10, FlowJo LLC).

#### EC panel

Equal number of cells were then blocked with 2µl FC-Block (Mouse BD Fc Block™) and 2.5µl rat serum (10710C, Thermo fisher scientific) per 100µl cell suspension for 10 minutes. After blocking cell were stained with FITC-conjugated Ter119 (116206, Biolegend), rabbit anti Ephrin-B2 (ab131536, abcam), eFluor 660- conjugated anti Endomucin (50-5851-82, Thermo fisher scientific, clone V.7C7), BV510-conjugatetd CD45 (103137, Biolegend, clone 30-F11) and PE-Cy7-conjugated CD31 (102418, Biolegend, clone 390) for 30 minutes on ice. After washing two times with cell stain buffer, secondary antibody (BV421-conjugated donkey anti-rabbit (406410, Biolegend) were added in brilliant stain buffer (BD) and 2.5µl donkey serum (017-000-121, Dianova) and incubated 30 minutes on ice. After a further washing step with cell stain buffer (420201, BioLegend), 7AAD viability staining solution (420404, Biolegend) was added and cells were acquired on BD FACS Canto II flow cytometer and analysed using FlowJo (Version 10, FlowJo LLC).

### Heart failure patients

Bone marrow aspirates were obtained from healthy volunteers without any evidence of coronary artery disease in their history (N=8, median age 31 years [IQR 27-37]) as well as from patients suffering from post-infarct heart failure with severely reduced left ventricular function, undergoing intracoronary infusion of bone marrow mononuclear cells within the REPEAT trial (Repetitive Progenitor Cell Therapy in Advanced Chronic Heart Failure; NCT 01693042, N=19). All controls and patients provided written informed consent. The ethics review board of the Goethe University in Frankfurt, Germany, approved the protocols (Approval No. 160/15 for healthy controls and the study consent for REPEAT trial)^1^. The study complies with the Declaration of Helsinki. Patients were eligible for inclusion into the study if they had stable post-MI HF symptoms New York Heart Association (NYHA) classification of at least II, had a previous successfully revascularized myocardial infarction at least 3 months before bone marrow aspiration, and had a well-demarcated region of left ventricular dysfunction on echocardiography. Exclusion criteria were the presence of acutely decompensated heart failure with NYHA class IV, an acute ischemic event within 3 months prior to inclusion, a history of severe chronic diseases, documented cancer within the preceding 5 years, or unwillingness to participate.

### Human bone marrow flow cytometry

Lineage negative and CD31 positive BMCs were isolated using first immunomagnetic Lineage Cell Depletion Kit (130-092-211, Miltenyi Biotec) followed by positive selection with CD31 MicroBead Kit (130-091-935, Miltenyi Biotec) using a magnetic cell separation device (QuadroMACS Separator;130-090-976, Miltenyi Biotec). Briefly, BMC suspensions were incubated for 10 minutes at 4°C with Biotin-Antibody Cocktail, washed with MACS Buffer, and then incubated for 15 minutes with anti-Biotin MicroBeads. After washing, cells were separated on LS columns (130-042- 401, Miltenyi Biotec) for positive and negative selection, respectively, according to the manufacturer’s instructions. After counting lineage negative cells, a second separating step with CD31 followed. Lineage negative cells were incubated with FcR Blocking Reagent and CD31 MicroBeads for 15 minutes at 4°C. After washing, cells were separated on LS columns (130-042-401, Miltenyi Biotec) for positive and negative selection, respectively, according to the manufacturer’s instructions. Lineage and CD31 expression of sorted fractions was checked by FACS-analysis with BV421-conjugated CD31 (303124, Biolegend, clone WM59) antibody and FITC-conjugated Lineage cocktail 4 (562722, BD, clones RPA-2.10, HIT3a, RPA-T4, M- T701, HIT8a, B159, GA-R2 (HIR2)).

100µl bone marrow were blocked with 2µl Fc Receptor Blocking Solution (Human TruStain FcX™, (422301, Biolegend) for 10 minutes at room temperature. BV-421- conjugated CD31 (303124, Biolegend, clone WM59), APC-Cy7-conjugated CD45 (368516, Biolegend, clone 2D1), FITC-conjugated Lineage cocktail 4 (562722, BD, clones RPA-2.10, HIT3a, RPA-T4, M-T701, HIT8a, B159, GA-R2 (HIR2)), APC-conjugated AC133 (130-090-826, Miltenyi Biotec), PE-Cy7-conjugated CD34 (343516, Biolegend, clone 581) and biotin-conjugated Endomucin (ab45772, abcam, clone TX18) were added to the bone marrow and incubated for 20 minutes. After staining BM was washed two times with cell stain buffer followed by a second incubation step with PE-conjugated Streptavidin (405204, Biolegend) for 20 minutes. Erythrocytes were lysed with 1x RBC lysis buffer (420301, Biolegend) for 10 minutes and washed two times with cell stain buffer. After adding 7AAD viability staining solution (420404, Biolegend), cells were measured on BD FACS Canto II flow cytometer and analysed using FlowJo (Version 10, FlowJo LLC).

### Human bone marrow immunocytochemistry

Human bone marrow endothelial cells were MACS-enriched from freshly drawn BM aspirates (in parallel from a healthy control and from a HF patient). The cell suspension was adjusted to 50,000 cells/100µl. 200µl of cell suspension (100.000 cells) were transferred into a Cytofunnel chamber and centrifuged for 5 minutes at 500xg, to allow for complete fluid absorption. Air-dried samples were fixed with 4% PFA, washed with PBS and permeabilized with 0.1% Triton X in PBS for 15 minutes at room temperature. After blocking, samples were incubated with primary antibody overnight at 4°C, followed by 3 wash steps (0.05% Triton X in PBS) and incubation with the secondary antibody for 1h at room temperature.

#### Primary antibodies

Mouse anti-c-Myc (MA1-980, Thermofisher) and Biotin anti-IL-1β (511703, Biolegend).

#### Secondary antibodies

Goat anti-mouse Alexa Fluor 555 (A-21425, Life Technologies) and Streptavidin Alexa Fluor 555 (S21381, Life Technologies).

### Cryo-section immunostaining

Freshly dissected bone tissues were fixed in ice-cold 4% paraformaldehyde solution for 4 hours. After PBS washing, decalcification was carried out with 10% EDTA/Tris/HCL pH=7.0 (15575-038, Invitrogen) at 4°C with constant shaking for 48 hours and decalcified bones were immersed into 20% sucrose (S0389, Sigma) and 1% polyvinylpyrrolidone (PVP) (P5288, Sigma) solution for 24 h. Finally, the tissues were embedded and frozen in 8% gelatin (porcine) (G1890, Sigma) in presence of 15% sucrose and 1% PVP. Cryosections were produced at a Leica CM3050S cryostat Cryosection were stored at −20°C. Before starting the staining procedure, they were allowed to acquired room temperature. Slides were then rehydrated in PBS and permeabilized with PBS containing 0.3% Triton X-100 by washing them three times 10 minutes. Tissue was blocked for one hour in freshly prepared blocking solution (3% BSA, 0.1% Triton X 100 and 0.5% donkey serum in PBS). Primary antibodies were incubated overnight at 4°C in blocking solution. The consecutive day, the primary antibodies were washed four times 5 minutes in PBS. Secondary antibodies were incubated for one hour at room temperature in PBS containing 5% BSA. Immunostainings were imaged in a Leica SP8 confocal inverted microscope.

#### Primary antibodies

Goat anti CD31 Alexa Fluor 488 conjugated (FAB3628G, R&D), rat anti Endomucin (sc-65495, Santa Cruz), rat anti CD41 Alexa Fluor 647 conjugated (133934, Biolegend), and mouse anti IL1β (12242, Cell Signalling).

#### Secondary antibodies

Donkey anti rat Alexa Fluor 594 (A21209, Life Technologies) and donkey anti mouse Alexa Fluor 647 (A31571, Life Technologies).

Stainings were quantified using Volocity Software 6.5 (Quorum Technologies)

### Bone RNA isolation and RT-qPCR

Bone RNA isolation was performed with Direct-zol RNA Miniprep Kits from (Zymoresearch, R2051) according to manufacturer protocol. RNA concentration was measured by nanodrop (Thermo). For cDNA synthesis, 1µg RNA was reverse-transcribed using M-MLV reverse transcriptase (Invitrogen, 28025-021). Quantitative PCR was performed with TaqMan fast Mastermix (435204, Applied Biosystems) and the following Taqman probes: IL1b (Mm00434228_m1) and P0 (Mm00725448_s1).

### Single cell RNA sequencing library preparation

Cellular suspensions were loaded on a 10X Chromium Controller (10X Genomics) according to manufacturer’s protocol based on the 10X Genomics proprietary technology. Single-cell RNA-Seq libraries were prepared using Chromium Single Cell 3′ Reagent Kit v2 (10X Genomics) according to manufacturer’s protocol. Briefly, the initial step consisted in performing an emulsion capture where individual cells were isolated into droplets together with gel beads coated with unique primers bearing 10X cell barcodes, UMI (unique molecular identifiers) and poly(dT) sequences. Reverse transcription reactions were engaged to generate barcoded full-length cDNA followed by the disruption of emulsions using the recovery agent and cDNA clean up with DynaBeads MyOne Silane Beads (Thermo Fisher Scientific). Bulk cDNA was amplified using a Biometra Thermocycler TProfessional Basic Gradient with 96-Well Sample Block (98°C for 3 minutes; cycled 14×: 98°C for 15s, 67°C for 20s, and 72°C for 1 minute; 72°C for 1 minute; held at 4°C). Amplified cDNA product was cleaned with the SPRIselect Reagent Kit (Beckman Coulter). Indexed sequencing libraries were constructed using the reagents from the Chromium Single Cell 3′ v2 Reagent Kit, as follows: fragmentation, end repair and A-tailing; size selection with SPRIselect; adaptor ligation; post-ligation cleanup with SPRIselect; sample index PCR and cleanup with SPRI select beads. Library quantification and quality assessment was performed using Bioanalyzer Agilent 2100 using a High Sensitivity DNA chip (Agilent Genomics). Indexed libraries were equimolarly pooled and sequenced on two Illumina HiSeq4000 using paired-end 26×98bp as sequencing mode.

### Single cell RNA sequencing data analyses

Single-cell expression data were processed using the Cell Ranger Single Cell Software Suite (v2.1.1) to perform quality control, sample de-multiplexing, barcode processing, and single-cell 3′ gene counting^2^. Sequencing reads were aligned to the human reference genome GRCh38 using the Cell Ranger suite with default parameters. Dimensional reduction analysis was performed in Seurat (v2) R (v3.6)^3^. Monocle (v2.6.0) was applied for differential expression analysis and the generation of cell trajectories^4^. The gene-cell-barcode matrix of the samples was log-transformed and filtered based on the number of genes detected per cell (any cell with less than 200 genes per cell or greater than 10% mitochondrial content was filtered out). Regression in gene expression was performed based on the number of unique molecular identifiers (UMI). PCA was run on the normalized gene-barcode matrix. Barnes-hut approximation to t-SNE was then performed on principal components to visualize cells in a two-dimensional space^5^. This graph-based clustering method relies on a clustering algorithm based on shared nearest neighbour (SNN) modularity optimization. Differential transcriptional profiles by cluster were generated in Seurat with associated gene ontology terms derived from the functional annotation tool DAVID Bioinformatics Resources 6.7 (NIAID/NIH, https://david.ncifcrf.gov/summary.jsp). Cell annotation was then performed by assessing relative expression of standard endothelial and hematopoietic markers. Cell fate trajectory analysis (Monocle) was utilized on an EMCN enriched cluster of cells to identify unique attributes between heart failure and healthy patients’ cells.

## Data availability

We will provide scRNA-seq data at the Gene expression Omnibus once the paper is accepted.

## Statistics

GraphPad’s 8 (Prism) software was used for statistical analysis of all experiments but scRNA-seq. Shapiro-Wilk normality test was used to test data before comparison. Unpaired, two-tailed Student’s t-test and Mann-Whitney test were used for comparison between two groups. For multiple comparisons, one-way ANOVA with Dunnet’s multiple comparisons test was used. Data is presented in scatter plots with mean ± standard error of mean (s.e.m). Differences were considered statistically significant at p<0.05.

